# Midfrontal theta is associated with errors, but no evidence for a link with error-related memory

**DOI:** 10.1101/2022.05.23.493097

**Authors:** Xiaochen Y. Zheng, Syanah C. Wynn

## Abstract

Midfrontal theta is widely observed in situations with increased demand for cognitive control, such as monitoring response errors. It also plays an important role in the cognitive control involved in memory, supporting processes like the binding of single items into a memory representation or encoding contextual information. In the current study, we explored the link between midfrontal theta and error-related memory. To this end, we recorded EEG from 31 participants while they performed a modified Flanker task. Their memory for the errors made during the task was assessed after each experimental block, and its relationship with error-related midfrontal theta effects was investigated. We have replicated the error-related increase in midfrontal theta power, reported in previous literature. However, this error-related theta effect could not predict subsequent memory of the committed errors. Our findings add to a growing literature on the prefrontal cortex-guided control process in error monitoring and episodic memory.

## Introduction

When we make a mistake, our brain sets in motion processes to prevent us from making additional mistakes. To learn from our errors, we need to monitor them and adjust our behavior accordingly. This happens on different timescales, in the milliseconds following an error, the brain needs to identify the event as an error to adjust immediate behavior accordingly. For instance, when accidentally hitting the gas pedal instead of the brake, our brain needs to quickly send a signal to our feet to initiate a motor response to correct this. In addition, we need to be able to remember the errors we made in the past to prevent them from happening again. The next time you step in a car you will remember the error you made hours, days, or weeks ago and you will be mindful of not making the same mistake again. Therefore, our brain needs the joint efforts of error-related and memory-related processes for us to learn from our mistakes in daily life.

The detection of an error is reflected by midfrontal theta oscillations (4-7 Hz), recorded from electroencephalography (EEG) channels over the medial prefrontal cortex (PFC; Cavanagh & Frank, 2014; Cohen, 2011; Fusco et al., 2018). Neural oscillations are thought to be important for transient brain computations needed for cognitive functions, and theta appears to coordinate processes needed for post-error cognitive control (Bonnefond, Kastner, & Jensen, 2017; Duprez, Gulbinaite, & Cohen, 2020; Fries, 2005; Jensen, Gips, Bergmann, & Bonnefond, 2014). For instance, when applied externally through transcranial alternating current stimulation (tACS), midfrontal theta modulated behavioral adjustments following errors in a Flanker task (Fusco et al., 2018). It is thought that midfrontal theta originates from the anterior cingulate cortex (ACC) and is cognitively associated with the monitoring of the errors (Botvinick, Cohen, & Carter, 2004; Cavanagh, Cohen, & Allen, 2009; Chevalier, Hadley, & Balthrop, 2021; Cohen, 2011). The increase in midfrontal theta may signal an increased need for top-down control to adjust behavior, recruiting the PFC to prevent subsequent errors (Cavanagh, Zambrano-Vazquez, & Allen, 2012; Cohen, 2011; Kerns, 2006). Given this involvement of midfrontal theta milliseconds after an error is made, it would be of interest to know if theta can also predict subsequent behavioral change in a longer time frame, such as memory for errors which could influence decisions at a later point in time.

As was the case for error processing, midfrontal theta also plays an important role in the cognitive control involved in episodic memory. PFC-guided top-down control is needed for the encoding and retrieval of target information, resolving competition with irrelevant information (Badre & Wagner, 2007; Blumenfeld, Parks, Yonelinas, & Ranganath, 2011; Blumenfeld & Ranganath, 2007; Nyhus & Badre, 2015). Successful memory processing therefore requires communication between the PFC and other relevant brain areas, like the medial temporal lobe (MTL), which is thought to occur through theta oscillations (Lin et al., 2017; Nyhus & Badre, 2015; Nyhus & Curran, 2010; Rutishauser, Ross, Mamelak, & Schuman, 2010; Staudigl & Hanslmayr, 2013; Sweeney-Reed et al., 2016; Wynn, Daselaar, Kessels, & Schutter, 2019; Wynn, Kessels, & Schutter, 2020). In concordance, EEG studies have shown that memory-related theta might mediate PFC-guided control processes needed for task-relevant encoding (Cavanagh & Frank, 2014; Nyhus & Badre, 2015; Nyhus& Curran, 2010). Midfrontal theta may support processes that are needed for episodic memory, like the binding of single items into a memory representation or encoding contextual information (Hsieh & Ranganath, 2014). Therefore, midfrontal theta appears to play a supporting role in both error monitoring and memory encoding.

In the current study, we aimed to bridge the error- and memory-related literature by exploring the link between midfrontal theta and error-related memory. Our participants performed a modified Flanker task (Eriksen & Eriksen, 1974), while we assessed their error memory after each experimental block. We hypothesized that midfrontal theta reflects the online detection of the errors and will predict participants’ memory of the errors they have made.

## Methods

### Participants

A total of 31 healthy right-handed adults participated in this study, recruited through the Radboud Research Participation System. All had normal or corrected-to-normal vision, were native Dutch speakers, and were free from any self-reported neurological or psychiatric conditions. All participants received course credit or monetary compensation. Of these 31 participants, three participants were excluded from the analyses reported here, due to limited number of trials left after artifact rejection (N=1) or data acquisition issues (N=2) that rendered the data unusable for data analyses reported here. This results in a total 28 participants (16 females, *M*_age_ = 22.43, *SD*_age_ = 3.86) reported in the current analyses. One additional participant without working memory (WM) measures was excluded from the correspondent analysis. The study was approved by the local ethics committee of the Faculty of Social Sciences of the Radboud University Nijmegen.

### Procedure

All participants received written information prior to participation, but remained naive regarding the aim of the study. Upon arrival at the laboratory, all participants were screened for eligibility to participate in EEG studies and provided written informed consent.

#### Working memory task

Prior to the Flanker task, a computerized version of the digit span task from the Weschler Adult Intelligence Scale fourth edition (WAIS-IV; Kreutzer, DeLuca, & Caplan, 2011) was used as a measurement of WM. The digit span task consisted of three conditions (forward, backward, and sequencing) and the order of these was kept consistent across participants. During all conditions, a single digit (1-9) was presented centrally on the screen for 1000 ms, followed by a 300 ms inter-stimulus interval. Digit presentation and recording of responses were attained using PsychoPy (v1.80; Peirce et al., 2019) on a Windows PC. For each condition, every trial consisted of two series of digits, which increased by one digit on every trial (e.g., first trial: 3-5 and 8-4; second trial: 9-5-2 and 1-7-6). In the forward condition, participants were asked to reproduce the digits in the same order as previously presented after each series (e.g., first trial: 3-5 and 8-4; second trial: 9-5-2 and 1-7-6). In the backward condition, they were asked to reproduce the series in the reversed order (e.g., first trial: 5-3 and 4-8; second trial: 2-5-9 and 6-7-1). In the sequencing condition, participants had to recall the digits in ascending order (e.g., first trial: 3-5 and 4-8; second trial: 2-5-9 and 1-6-7). Participants responded by typing the digit sequence on a keyboard. Participants were able to alter their response up to the moment of confirmation, which was operationalized by pressing the enter key. The task was aborted when a participant was not able to respond correctly in both two series in a single trial. The total number of correct responses was used as their WM score. The maximal score that could possibly be obtained was 16 for all three conditions.

#### Flanker task

Thereafter, participants performed a modified Flanker task (Eriksen & Eriksen, 1974), where they were required to give a speeded response to a central target arrow, while ignoring congruent (“≫>≫” or “≪<≪”) or incongruent (“≫<≫” or “≪>≪”) Flanker arrows. Half of the trials were congruent and half were incongruent. Participants responded to the target arrows by pressing the “D” key (left arrows) and the “H” key (right arrows) on a keyboard, with their left and right index fingers, respectively. Stimuli were presented on a grey background for 200 ms, with stimulus onset asynchronies randomly selected from a uniform distribution with a mean of 1550 ms and varying between 1400 and 1700 ms with 50 ms increments. During the intertrial interval, a white cross was centrally presented and participants were instructed to keep fixation on the cross.

To elicit a sufficient number of errors, we trained the participants to respond within a time limit prior to the main task. The time limit was computed dynamically across the training and calibrated individually for each participant (based on the 70 percentile of previous ten trials). If participants failed to respond within a given time limit, the fixation cross became red as a warning. The speed training consisted of 100 trials.

The main task that followed, was divided into six blocks of 100 trials. Participants were not given feedback on their performance during the main task. At the end of each block, participants were instructed to recall their task performance. Specifically, after each block, participants were asked to recall the number of errors made in the preceding block and rate their confidence in this judgment. To minimize any influence on task strategy, we also asked them to recall the number of times they felt their response was too slow in the preceding block. The numerical responses were given with the numbers on the keyboard, and the confidence ratings were submitted by clicking with a mouse on a visual analog scale, ranging from “completely not sure” to “completely sure”.

#### EEG acquisition

EEG signals were recorded and amplified with a BioSemi ActiveTwo system (BioSemi B.V., Amsterdam) from 32 Ag/AgCl-tipped electrodes, conforming to the International 10-20 System. The EEG signal was digitized at a sampling rate of 1024 Hz. Reference electrodes were placed bilateral on the mastoids, and bipolar electro-oculogram recordings were obtained from electrodes placed 1 cm lateral of the outer canthi, and above and below the left eye. Each active electrode was measured online with respect to a Common Mode Sense (CMS) active electrode. BioSemi uses a combination of a CMS electrode and a Driven Right Leg (DRL) passive electrode to ensure that the CMS electrode stays as close as possible to the reference voltage at the analogue-to-digital converter.

### Data Analysis

Data analyses were performed in MATLAB (v2021a; MathWorks Inc., Natrick MA) in combination with Fieldtrip toolbox (v20200128; Oostenveld, Fries, Maris, & Schoffelen, 2011), and in R (Version 4.0.2; R Core Team, 2013).

#### EEG pre-processing

Continuous data were first re-referenced to linked-mastoid references and then band-pass filtered with a low cut-off of 0.1 Hz and a high cut-off of 30 Hz. We then segmented the data into epochs from 500 ms before to 1400 ms after stimulus onset. Trials with atypical artifacts (e.g., jumps and drifts) as well as bad channels (less than 0.3%) were rejected by visual inspection; EOG artifacts (eye blinks and saccades) were removed using independent component analysis. After ICA, we reconstruct the earlier rejected channels by a weighted average of the data from neighboring channels of the same participant. The data were further segmented into epochs from 500 ms before to 800 ms after response onset. In an additional round of visual inspection, trials with remaining artifacts were removed.

#### Time-frequency analysis

To determine the time window that was sensitive to error-related processes, a cluster-based permutation on the time-frequency representations (TFRs) was performed (Oostenveld et al., 2011). First, spectral power was extracted using Fourier analysis with sliding time windows and the application of a Hanning taper. Data was padded to two seconds and frequencies were assessed from 1 to 30 Hz, with a fixed 500 ms time window length for each frequency. Then TFRs of all error and correct trials were pooled together across participants. Based on the midfrontal theta literature (e.g., Cavanagh & Frank, 2014; Cohen, 2011; Fusco et al., 2018), we restrict all our analysis to a selection of frontal midline channels (‘FC1’, ‘FC2’, ‘Fz’, ‘Cz’) and the theta frequency band (4 - 7 Hz). We averaged the data in the channels and frequencies of interest in this analysis. To explore the effect in the time domain, all time points (i.e., −500 to 800 ms, time locked to the response onset) were included in the analysis. For every sample, the error and correct conditions were compared by means of a t-value. All samples with an α-value smaller than .05 were selected and clustered. The corresponding cluster-level statistics were calculated by taking the sum of the *t*-values within each cluster. The largest cluster-level statistic was used as the observed cluster-based test statistic. The cluster-based test statistic distribution was approximated utilizing the Monte Carlo method with 10,000 random partitions. The proportion of random partitions that resulted in a larger test statistic than the observed one (the Monte Carlo significance probability) was compared to the critical α-value of .05 (two-sided). If the Monte Carlo significance probability was smaller than .05, the data in the error and correct conditions were considered significantly different.

#### Trial-by-trial frequency analysis

To be able to look at theta power over trials, the data was first re-segmented to the 0-500 ms time window, relative to response onset and then padded to 1 second. This time window was chosen based on the results of the TFR analysis. For every trial, Fourier analysis was used to obtain the spectral decomposition of this data, using a Hanning taper. This gave the average theta power over the frontocentral channels in the 0-500 ms time window after response onset for each trial.

#### Mixed-effect models

The error memory performance per block was calculated as:

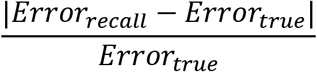

Where *Error_recall_* is the number of errors the participants remembered and *Error_true_* the actual number of errors made.

For the error-related midfrontal theta effect, we used the data from the trial-by-trial frequency analysis. Per participant and per block, the average theta power difference between error and correct trials was calculated as:

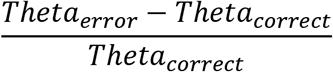

We used linear mixed-effects models utilizing the lme4 package (Version 1.1.27; Bates et al., 2011). Errors and RTs were modeled trial by trial. The errors were analyzed using generalized linear mixed-effects models (binomial family), as a function of Flanker congruency, post-error status (i.e., whether the current trial followed an error trial), WM score, and block number. Flanker congruency and post-error status were included as random slopes for participants. RT data were log-transferred to account for its right-skewed distribution. We additionally included accuracy as a fixed effect as well as a random slope for participants when modeling the RTs.

Error memory performance was modeled by block. We modeled error memory as a function of the error-related midfrontal theta effect, error rate, WM score, and block number. Error rates were included as a random slope for participants.

All models used in the analyses are provided in Supplementary material A.

## Results

### Flanker task performance and working memory

Participants’ performance on the Flanker task is shown in *Figure 1*. In line with the literature, participants showed a congruency effect; they were slower (β = 0.15, *SE* = 0.01, *t* = 9.94, *p* < .001) and made more errors (β = 2.38, *SE* = 0.19, *z* = 12.57, *p* < .001) in the incongruent than the congruent condition. In addition, they were faster when making an erroneous response as compared to a correct one (β = − 0.20, *SE* = 0.02, *t* = −11.21, *p* < .001). We observed post-error slowing (β = 0.01, *SE* = 0.005, *t* = 2.82, *p* = .009), but no post-error accuracy change (β = 0.04, *SE* = 0.08, *z* = 0.47, *p* = .64). Over time, participants got faster (β = −0.02, *SE* = 0.003, *t* = −5.56, *p* < .001), but their accuracy remained the same (β = −0.03, *SE* = 0.06, *z* = −0.53, *p* = .60).

**Figure 1.**
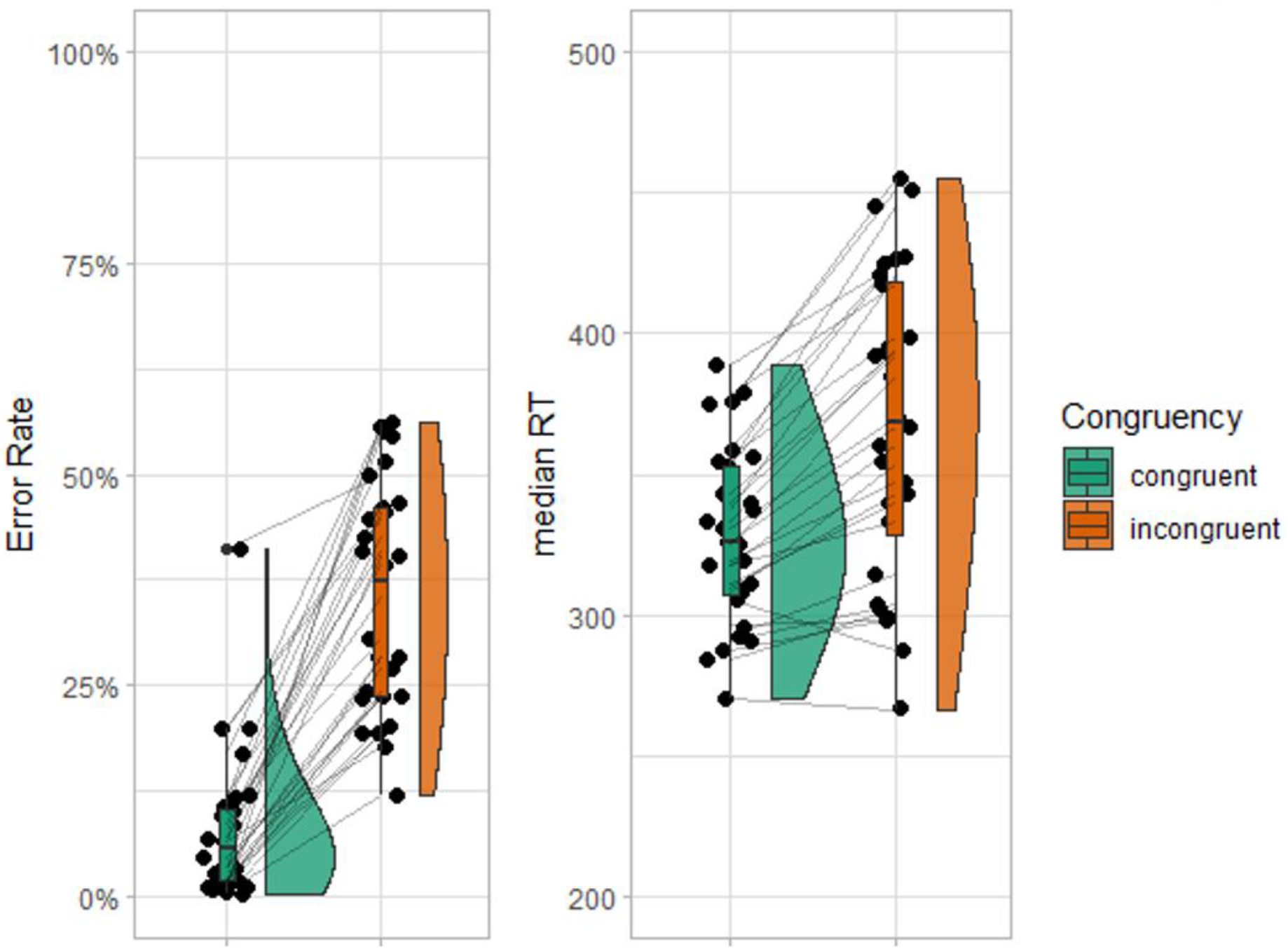
Raincloud plots of error rate (left panel) and median reaction time (RT, right panel) as a function of trial congruency. The outer shapes represent the distribution of the data over participants, the thick horizontal line inside the box indicates the group median, and the bottom and top of the box indicate the group-level first and third quartiles of each condition. Each dot represents one participant, the thin lines in between connect the same participant’s data for different conditions.

Participants’ working memory performance was quantified as the total number of correct responses on the digit span task. Participant had an average total WM score of 32 (*M* = 31.85, *SD* 5.50) over the three subtasks (*M*_forward_ 9.93, *SD*_forward_ 2.09; *M*_backward_ 10.81, *SD*_backward_ 2.40; *M*_sequencing_ = 11.11, *SD*_sequencing_ = 2.50). Their WM score could not predict their performance on the Flanker task (RT: β < 0.001, *SE* = 0.002, *t* = 0.32, *p* = .75; accuracy: β = 0.01, *SE* = 0.02, *z* = 0.66, *p* = .51).

### Midfrontal theta is modulated by Flanker errors

We explored whether theta power was modulated by Flanker performance, as suggested by the literature (e.g., Nigbur, Ivanova, & Sturmer, 2011), and inspect the temporal nature of this effect. When we look at *Figure 2*, comparing error and correct trials, there appears to be an increase in theta power over midfrontal channels in the first 250 ms after an error is made. This observation was tested by a cluster-based permutation analysis, which revealed that theta power increased significantly following an erroneous, compared to a correct response (*p* < .001). This effect was most pronounced between −31 ms and 545 ms relative to response onset. Based on visual inspection of this effect, further analyses in the manuscript were restricted to 0-500 ms.

**Figure 2.**
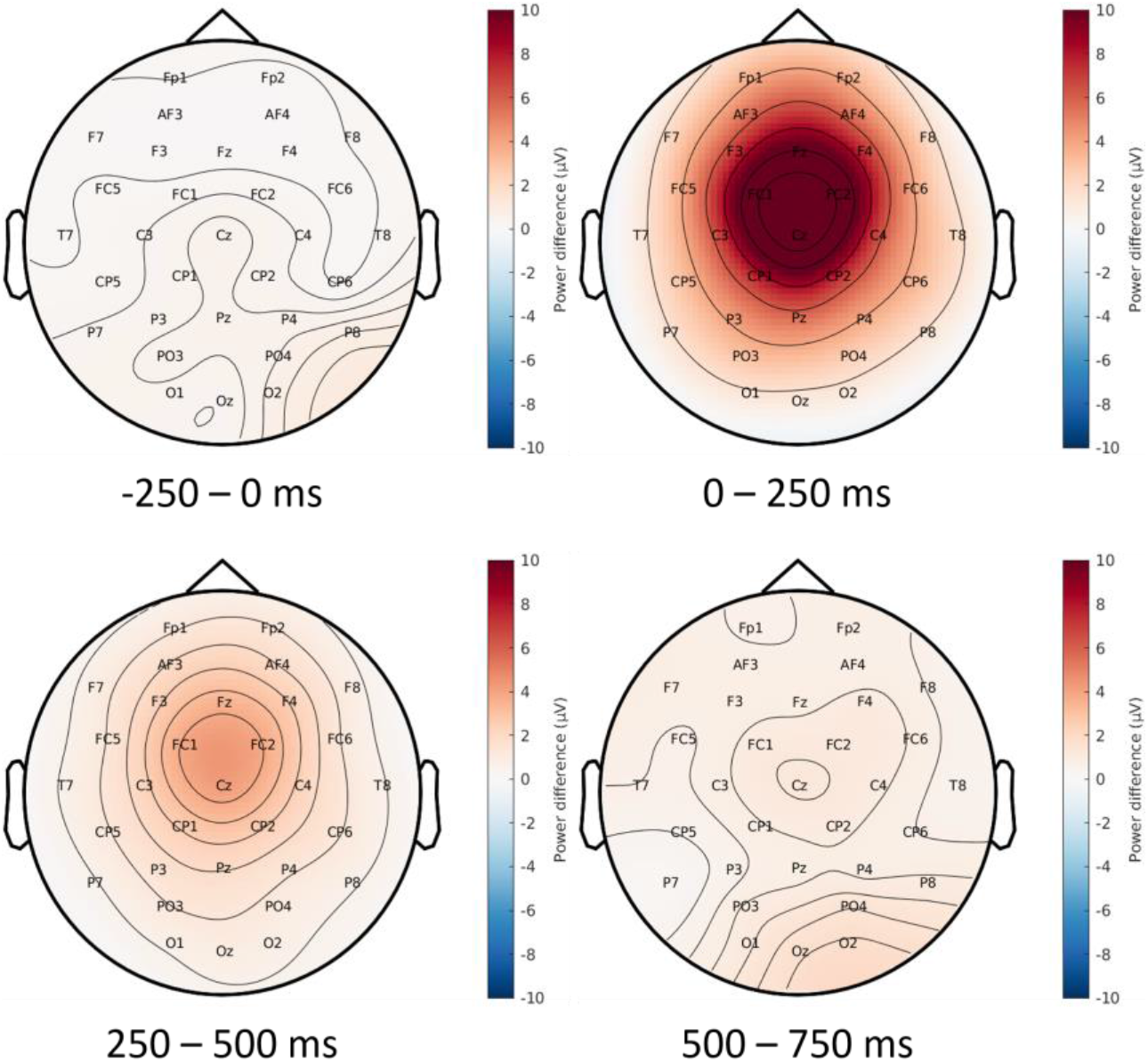
The theta power difference between error and correct trials over time. All time windows are relative to response onset.

### Error-related midfrontal theta cannot predict error memory performance

Participant’ ability to remember their errors across blocks is visualized in Figure 3. As can be seen in this figure, error memory performance did not change over time (β = 0.09, *SE* = 0.06, *t* = 1.56, *p* = .12).

**Figure 3.**
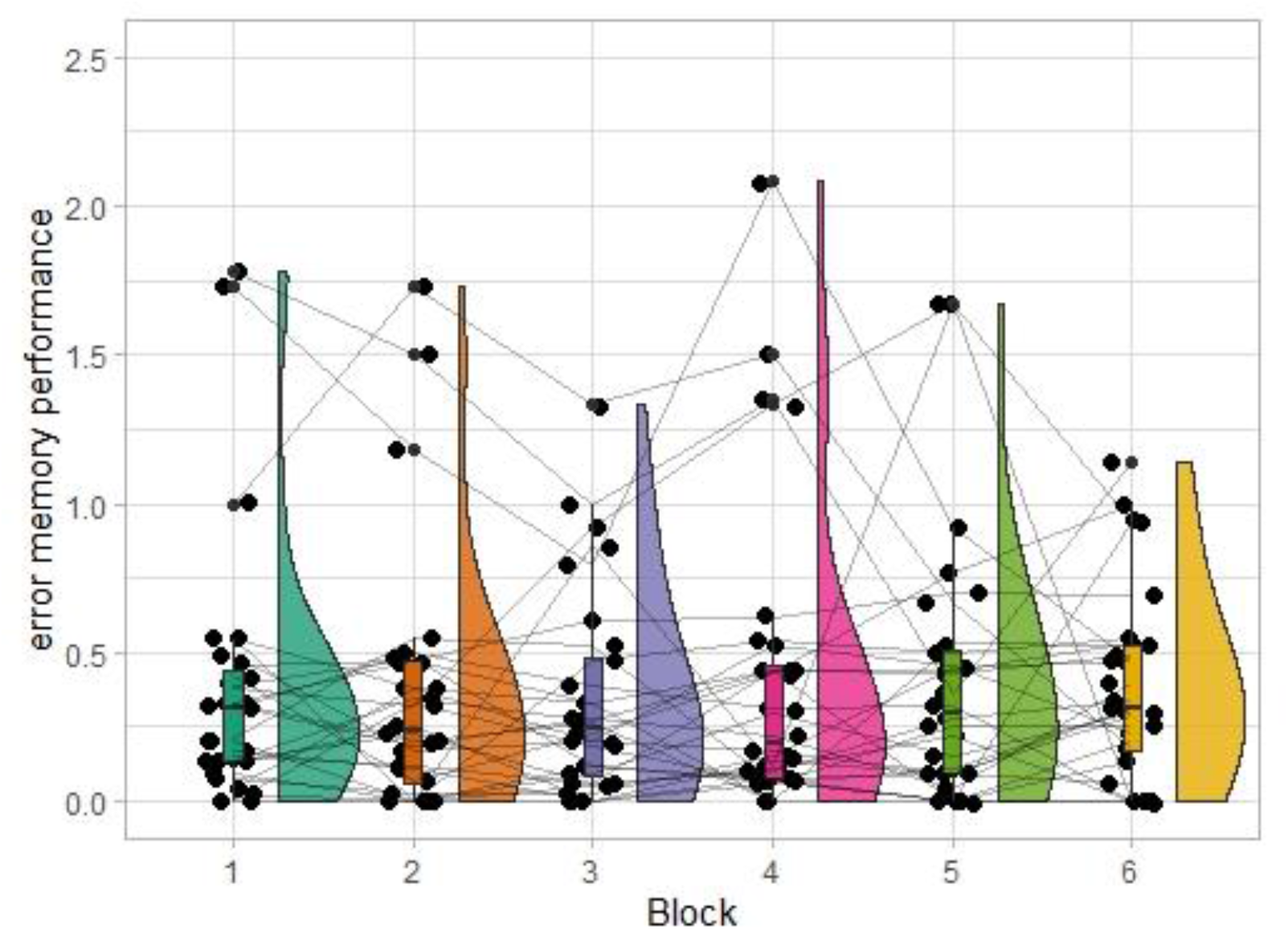
Raincloud plots of participants’ error memory performance as a function of block numbers. The smaller the value, the more accuracy participants were in recalling their errors. The outer shapes represent the distribution of the data over participants, the thick horizontal line inside the box indicates the group median, and the bottom and top of the box indicate the group-level first and third quartiles of each condition. Each dot represents one participant, the thin lines in between connect the same participant’s data for different blocks.

Figure 4 (left panel) shows the relationship between participants’ error-related theta effect during each Flanker block and their error memory performance after each block. In general, it appears that there is no clear relationship between this neural theta effect and the memory performance. In addition, there also seems to be very little consistency between participants as can be seen in three example participants in the right panel of Figure 4. This observation was tested by utilizing a linear mixed effects model. The model showed that error memory performance could not be predicted by the error-related midfrontal theta effect (β = 0.01, *SE* = 0.04, *t* = 0.31, *p* = .76). This indicated that contrary to our prediction, online modulation of error-related midfrontal theta was not predictive of later error memory.

**Figure 4.**
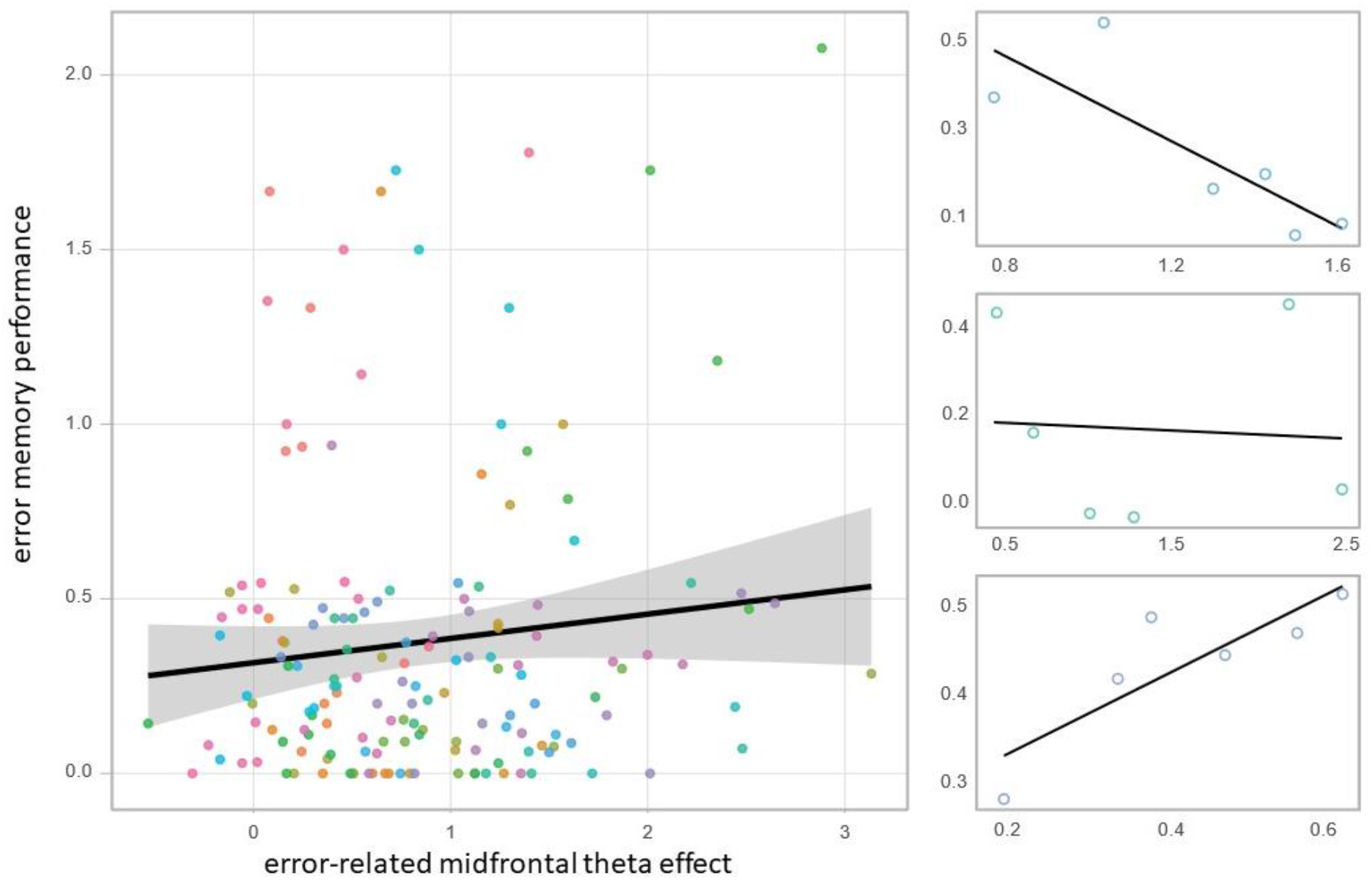
The relationship between the error-related midfrontal theta effect and error memory performance. Left panel: Each color codes for an individual participant and the corresponding regression line. The black regression line fits all the data points, and the gray area depicts the confidence interval. Right panel: three example participants. The color corresponds to the ones on the left panel.

We also accounted for participants’ WM score and their performance accuracy in the same model. Neither error rates (β = −0.35, *SE* = 0.63, *t* = −0.56, *p* = .58), nor participants’ WM score (β = −0.006, *SE* = 0.01, *t* = −0.57, *p* = .57) affected their error memory performance.

## Discussion

Midfrontal theta is enhanced in situations that call for more cognitive control (Cavanagh & Frank, 2014; Cavanagh et al., 2012). One of these instances is the occurrence of an error, where cognitive control is needed for subsequent behavioral adjustments (Cavanagh et al., 2012; Cohen, 2011; Fusco et al., 2018; Luu, Tucker, & Makeig, 2004). On the other hand, midfrontal theta also plays a crucial role in memory-related processes (Hsieh & Ranganath, 2014). For example, theta is higher during memory encoding for items that are subsequently recollected (Hanslmayr, Spitzer, & Bauml, 2009; Summerfield & Mangels, 2005; White et al., 2013). In the current study, we examined the link between the error-related midfrontal theta effect and participants’ memory of the errors they have made.

While participants were performing a modified Flanker task, we recorded their EEG activity and compared response-related theta power after erroneous and correct responses. Our results are in line with previous literature on the involvement of midfrontal theta in error processing (Cavanagh et al., 2012; Cohen, 2011; Luu et al., 2004; Nigbur et al., 2011). Like previous studies, we observed enhanced theta power following an error commission as compared to a correct response. This error-related theta effect was present mainly in the medial frontal scalp region and in the first 500 ms after a response was made. This theta effect likely reflects error detection and the signaling of post-error cognitive control (Bonnefond et al., 2017; Cavanagh & Frank, 2014; Duprez et al., 2020; Fries, 2005).

Does the detection of response errors also affect the memory recollection of these errors? We asked participants to indicate how many errors they remembered making in each experimental block and explored the relationship between their memory performance and the trial-by-trial brain oscillation. Our results provide no evidence that error memory can be predicted by the error-related theta effect, which seems to suggest a discrepancy between the error-related and memory-related control processes. In line with the idea that the same neural implementations can be driven by distinct neuronal computation principles (Buzsaki, Anastassiou, & Koch, 2012), it is plausible that error-related and memory-related midfrontal theta have different underlying mechanisms. For instance, midfrontal theta has long been viewed as the EEG signature of the unitary process of response conflict detection. However, it has been recently proposed that midfrontal theta reflects multiple uncorrelated processes, which give rise to comparable EEG compositions (Beldzik, Ullsperger, Domagalik, & Marek, 2022; Zuure, Hinkley, Tiesinga, Nagarajan, & Cohen, 2020). Therefore, it is plausible that even though error-related and memory-related control processes are both associated with “midfrontal theta”, the underlying neural mechanisms differ and are uncoupled.

An influential hypothesis is that theta oscillations may coordinate the timing of cognitive processes due to large-scale cross-frequency coupling (Duprez et al., 2020; Lisman & Jensen, 2013). Processes specific to post-error control may be linked to a specific midfrontal theta phase, while another cognitive process, like memory encoding could have a different preferential theta phase. Information arriving at specific phases would initiate multiple parallel processes in various brain regions. For instance, after error detection, communication in an extended neural network comprising of the PFC, medial temporal lobe and posterior parietal cortex might be required for subsequent memory and decision-making processes (Cohen, 2011; Thakral, Wang, & Rugg, 2017). In concordance, it has been proposed that theta oscillations mediate top-down control from the PFC to the hippocampus for selective encoding and retrieval of episodic memories (Nyhus & Curran, 2010). It is therefore a possibility that the initial increase in theta power after an error is made, is not predictive of subsequent error memory due to additional processing that occurs afterwards.

Several limitations of the study should be mentioned. First, contrary to most memory studies, our measure of error-related theta does not reflect a contrast between successful and unsuccessful encoding. It would be ideal if we could dissociate errors that have been successfully and unsuccessfully encoded during the task, which however was not possible in the current design. With state-of-art pattern analysis and decoding techniques of the neuronal data, future studies might be able to examine this further. Second, we quantified memory accuracy as the absolute difference between recalled errors and truly committed errors. This quantification does not differentiate between errors that were forgotten (misses) and correct trials misremembered as errors (false alarms). It could be that we found no evidence for a link between error-related theta and error memory due to the pooling of these errors. Since we could not directly differentiate between misses and false alarms, we used the error under- or overestimation as a proxy and explored its link with error-related theta. Nevertheless, our exploratory analysis (Supplementary Material B) shows no evidence that midfrontal theta can predict misses and false alarms.

To summarize, we have replicated the error-related midfrontal theta using a modified Flanker task. However, the error-related theta power increase cannot predict subsequent memory performance of committed errors. These findings add to a growing literature on the PFC function in cognitive control and memory process. Still, much remains to be explored on whether midfrontal theta, a seemingly distinctive neural signature of cognitive control, actually reflects multiple cognitive processes.

## Data Availability

Data is available at the Donders Repository (https://data.donders.ru.nl/). They will be shared publicly upon manuscript acceptance.

## Supplementary Materials

### Supplementary Material A: (generalized) linear mixed effect models

#### # for accuracy

glmer(accuracy ~ congruency + post_error + WM_sum + ordered(block) + (1 + congruency + post_error | participant), data = data_by_trial, family − ‘binomial”, control = glmerControl(optimizer =“bobyqa”))

#### # for RT

lmer(log(RT) ~ congruency + accuracy + post_error + WM_sum + ordered(Block) + (1 + congruency + accuracy + post_error | participant), data = data_by_trial, control = lmerControl(optimizer =“bobyqa”))

#### # for error memory performance

lmer(error_memory_abs ~ theta_effect + error_rate + WM_sum + ordered(block) + (1 + error_rate | participant), data = data_by_block, control = lmerControl(optimizer =“bobyqa”))

#### # additional analysis for error memory performance (over- vs. under-estimation)

glmer(error_memory_sign ~ theta_effect + error_rate + ordered(block) + (1 | participant), data = data_by_block, family =“binomial”, control = glmerControl(optimizer =“bobyqa”))

### Supplementary Material B: additional analysis

**Figure S1.**
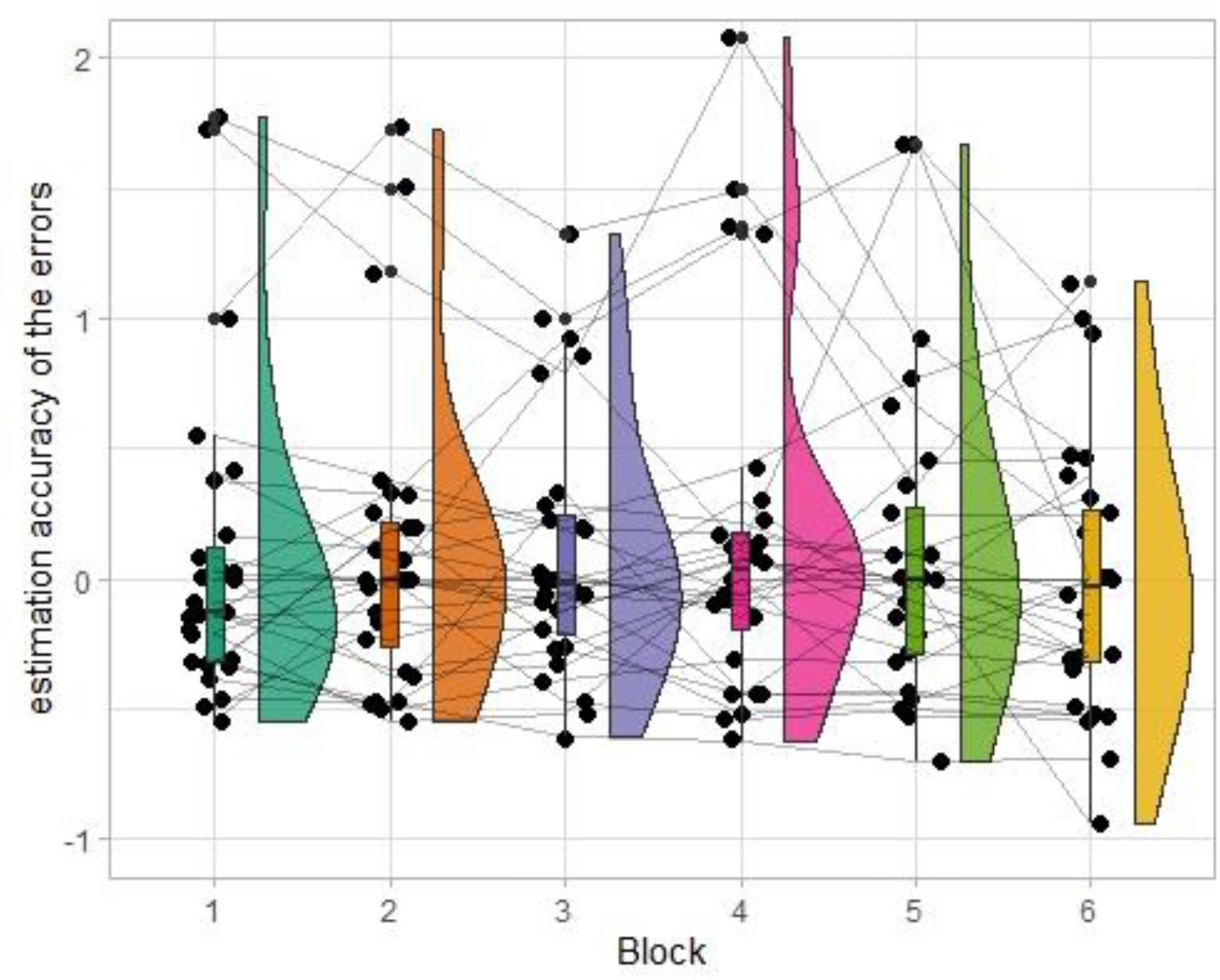
Raincloud plots of participants’ estimation memory performance, computed as:

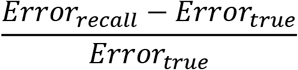 A positive value means overestimating, a negative value means underestimating. The outer shapes represent the distribution of the data over participants, the thick horizontal line inside the box indicates the group median, and the bottom and top of the box indicate the group-level first and third quartiles of each condition. Each dot represents one participant, the thin lines in between connect the same participant’s data for different blocks.

In general, participants tend to overestimate rather than underestimate the number of errors they made (Figure S1). However, this cannot be predicted by their error-related midfrontal theta effect (β = −0.34, *SE* = 0.42, *t* = −0.83, *p* = .41). Additionally, although participants tend to underestimate when they make more errors (β = −13.47, *SE* = 4.04, *t* = −3.33,*p*< .001), their tendency to over- or underestimate do not change over time (β = 0.09, *SE* = 0.53, *z* = 0.18, *p* = .86).

